# Specification of the germline by Nanos-dependent down-regulation of the somatic synMuvB transcription factor LIN-15B

**DOI:** 10.1101/163642

**Authors:** Chih-Yung S. Lee, Tu Lu, Geraldine Seydoux

## Abstract

The Nanos RNA-binding protein has been implicated in the specification of primordial germ cells (PGCs) in metazoans, but the underlying mechanisms remain poorly understood. We have profiled the transcriptome of PGCs lacking the *nanos* homologues *nos-1* and *nos-2* i*C. elegans*. *nos-1nos-2* PGCs fail to silence hundreds of genes normally expressed in oocytes and somatic cells, a phenotype reminiscent of PGCs lacking the repressive PRC2 complex. The *nos-1nos-2* phenotype depends on LIN-15B, a broadly expressed synMuvB class transcription factor known to antagonize PRC2 activity in somatic cells. LIN-15B is maternally-inherited by all embryonic cells and is down-regulated specifically in PGCs in a *nos-1nos-2*-dependent manner. Consistent with LIN-15B being a critical target of Nanos regulation, inactivation of maternal LIN-15B restores fertility to *nos-1nos-2* mutants. These studies demonstrate a central role for Nanos in reprogramming the transcriptome of PGCs away from an oocyte/somatic fate by down-regulating an antagonist of PRC2 activity.

## Introduction

In animals, formation of the germline begins during embryogenesis when a few cells (~30 in mice, 2 in *C. elegans*) become fated as primordial germ cells (PGCs) – the founder cells of the germline. PGC specification requires the activity of chromatin regulators that induce genome-wide changes in gene expression. For example, in mice, the transcriptional repressor BLIMP1 initiates PGC specification by blocking the expression of a mesodermal program active in neighboring somatic cells (Ohinata et al., 2005; Saitou et al., 2005). In *C. elegans*, the NSD methyltransferase MES-4 and the PRC2 complex (MES-2, 3 and 6) cooperate to place active and repressive histone marks on germline and somatic genes, respectively (Gaydos et al., 2012). Despite their critical roles during germ cell development, the MES and BLIMP1 regulators are not germline-specific factors and also function during the differentiation of somatic lineages (Cui et al., 2006; Gaydos et al., 2012; Seydoux and Braun, 2006). How the activities of these global regulators are modulated in germ cells to promote a germline-specific program is not well understood In *C. elegans*, genetic analyses have shown that MES activity is antagonized in somatic lineages by the synMuvB group of transcriptional regulators (Curran et al., 2009; Petrella et al., 2011; Unhavaithaya et al., 2002). Loss-of-function mutations in synMuvB genes cause ectopic activation of germline genes in intestinal cells and result in larval growth arrest at elevated temperatures (26஬). Inactivation of MES proteins suppresses the ectopic germline gene expression and restores viability to synMuvB mutants (Petrella et al., 2011). A similar antagonism has been uncovered in the adult germline between *mes-4* and the synMuvB gene *lin-54* (Tabuchi et al., 2013). The X chromosome is a major focus of MES repression in *C. elegans* germline. The X chromosome is silenced throughout germ cell development except in oocytes, which activate the transcription of many X-linked genes in preparation for embryogenesis (Kelly et al., 2002). *mes* mutants prematurely activate the transcription of somatic and X-linked genes in pre-gametic germ cells leading to germ cell death (Bender et al., 2006; Gaydos et al., 2012; Seelk et al., 2016). Reducing the function of the synMuvB transcription factor *lin-54* in *mes-4* mutant restores the expression of X-linked genes closer to wild-type levels (Tabuchi et al., 2013). Together, these genetic studies suggest that competition between the MES chromatin modifiers and the synMuvB class of transcription factors tunes X chromosome silencing and the ratio of soma/germline gene expression in somatic and germline tissues. How the balance of synMuvB/MES activities is initially set for each tissue, however, is not known.

The *C. elegans* PGCs arise early in embryogenesis from pluripotent progenitors (P blastomeres) that also generate somatic lineages. RNA polymerase II activity is repressed in the P lineage until the 100-cell stage when the last P blastomere P_4_ divides to generate Z2 and Z3, the two PGCs (Seydoux et al., 1996). RNA polymerase II becomes active in PGCs, but these cells remain relatively transcriptionally quiescent, and exhibit reduced levels of active chromatin marks compared to somatic cells throughout the remainder of embryogenesis (Kelly, 2014). Active marks and robust transcription return after hatching when the L1 larva begins to feed and the PGCs resume proliferation in the somatic gonad (Fukuyama et al., 2006; Kelly, 2014). The mechanisms that maintain PGC chromatin in a silenced state during embryogenesis are not known, but embryos lacking the *nanos* homologs *nos-1* and *nos-2* have been reported to display abnormally high levels of the H3meK4 mark in PGCs (Schaner et al., 2003). *nos-1nos-2* PGCs initiate proliferation prematurely during embryogenesis and die during the second larval stage (Subramaniam and Seydoux, 1999). Nanos proteins are broadly conserved across metazoans and have been shown to be required for PGC survival in several phyla, from insects to mammals (Asaoka-Taguchi et al., 1999; Beer and Draper, 2013; Deshpande et al., 1999; Lai et al., 2012; Tsuda et al., 2003). Nanos proteins are cytoplasmic and regulate gene expression post-transcriptionally by recruiting effector complexes that silence and degrade mRNAs in the cytoplasm. Six direct Nanos mRNA targets have been identified to date [Drosophila *hunchback, cyclin B and hid*(Asaoka-Taguchi et al., 1999; Dalby and Glover, 1993; Kadyrova et al., 2007; Murata and Wharton, 1995; Sato et al., 2007; Wreden et al., 1997), *Xenopus VegT* (Lai et al., 2012), and sea urchin *CNOT6* and *eEF1A* (Oulhen et al., 2017; Swartz et al., 2014)], but none of these targets are sufficient to explain how Nanos activity might affect PGC chromatin. In this study, we characterize the gene expression defects of PGCs lacking *nanos* activity in *C. elegans*. Our findings indicate that *nanos* activity is required to silence a maternal program active in oocytes and somatic embryonic cells. We identify the synMuvB transcription factor *lin-15B* as a critical target of Nanos regulation and demonstrate that down-regulation of maternal LIN-15B is essential to establish PRC2 dominance in PGCs.

## Results

### PGCs lacking *nos-1* and *nos-2* upregulate oogenic genes

*nos-2* is provided maternally and functions redundantly with zygotically expressed *nos-1* (Subramaniam and Seydoux, 1999). To generate large numbers of larvae lacking both *nos-1* and *nos-2* activities, we fed hermaphrodites homozygous for a deletion in *nos-1* [*nos-1(gv5)*] bacteria expressing *nos-2* dsRNA and collected their progeny at the L1 stage [hereafter designated *nos-1(gv5)nos-2(RNAi)* L1 larvae]. We used fluorescence-activated cell sorting (FACS) to isolate PGCs based on expression of the germ cell marker PGL-1::GFP and processed the sorted cells for RNA-seq (L1 PGCs). Two independent RNA-seq libraries (biological replicates) were analyzed for each genotype (wild-type and *nos-1(gv5)nos-2(RNAi)*) using Tophat 2.0.8 and Cufflinks 2.0.2 software (Trapnell et al., 2012). These analyses identified 461 under-expressed transcripts and 871 over-expressed transcripts in *nos-1(gv5)nos-2(RNAi)* L1 PGCs compared to wild-type (q >0.05, Figure 1A and Table S5 for list of miss-regulated genes). qRT-PCR of 11 genes confirmed the result of the RNA-seq analysis (Figure S1A).

**Figure 1.**
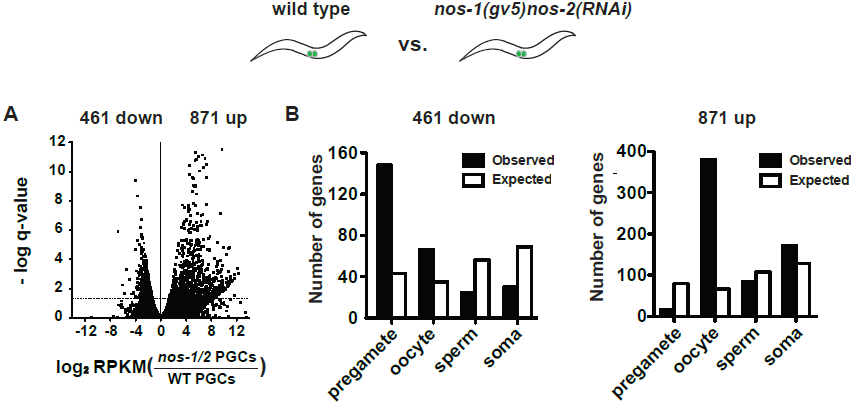
*nos-1nos-2* PGCs upregulate oogenic genes. Transcriptome comparison between PGCs isolated from wild-type and *nos-1(gv5)nos-2(RNAi)* L1 larvae using SMART-seq libraries (Materials and methods, See Figure S1B-C for results with Truseq libraries). (A) Volcano plot showing log2 fold-change in transcript abundance for each gene. The number of genes that were significantly up- or down-regulated in *nos-1(gv5)nos-2(RNAi)* PGCs are indicated. Dashed line marks the significance cutoff of q = 0.05 (Y axis) above which genes were counted as miss-expressed. (B) Bar graphs showing expected and observed number of genes (Y axis) in different expression categories (X axis). Genes were assigned to a particular expression category based on their preferential expression patterns as determined in (Gaydos et al., 2012; Ortiz et al., 2014) (Table S1). The lists are non-overlapping and include 2064 pregamete genes, 1688 oocyte genes, 2748 sperm genes, and 3239 somatic genes. Because genes were categorized based on their preferential gene expression pattern, genes on one list may also be expressed in other tissues. See Table S1 for complete gene lists. Pre-gamete genes are overrepresented among down-regulated genes and oocyte genes are overrepresented among up-regulated genes.

To determine the types of genes affected, we used published gene expression data (Gaydos et al., 2012; Meissner et al., 2009; Ortiz et al., 2014; Reinke et al., 2004; Wang et al., 2009) to generate non-overlapping lists of genes with preferential expression in pre-gametic germ cells, oocytes, sperm, or somatic cells (described and listed in Table S1).

We found that 32% (148/461) of under-expressed transcripts in *nos-1(gv5)nos-2(RNAi)* L1 PGCs correspond to genes expressed preferentially in pre-gametic germ cells (Figure 1B). These include *sygl-1,* a gene transcribed in germline stem cells in response to Notch signaling from the somatic gonad (Kershner et al., 2014). The *sygl-1* transcript was decreased by 4.7-fold in *nos-1(gv5)nos-2(RNAi)* PGCs. In contrast, over-expressed transcripts in *nos-1(gv5)nos-2(RNAi)* L1 PGCs correspond primarily to genes expressed in oocytes (380/ 871) (Figure 1B). These include *lin-41,* a master regulator of oocyte fate (Spike et al., 2014a; 2014b). The *lin-41* transcript was up-regulated by 5.1-fold in *nos-1(gv5)nos-2(RNAi)* PGCs. We conclude that *nos-1(gv5)nos-2(RNAi)* PGCs over-express oogenic genes and fail to activate pre-gametic genes normally expressed in PGCs.

### Turnover of maternal transcripts is delayed in PGCs lacking *nos-1* and *nos-2*

Oogenic transcripts in *nos-1(gv5)nos-2(RNAi)* L1 PGCs could correspond to maternal transcripts that failed to turnover during embryogenesis or to zygotic transcripts synthesized *de novo* in *nos-1(gv5)nos-2(RNAi)* PGCs. To distinguish between these possibilities, we isolated PGCs from embryos with fewer than 200 cells, at a time when PGCs are still mostly transcriptionally silent (EMB PGCs) (Schaner et al., 2003; Seydoux and Dunn, 1997). By comparing the EMB PGC transcriptome to a published oocyte transcriptome (Stoeckius et al., 2014), we observed an excellent correlation in relative transcript abundance between oocytes and EMB PGCs (Figure S2A). This observation suggests that many maternal mRNAs are maintained in the nascent germ lineage up to the 200-cell stage, as suggested earlier by *in situ* hybridization experiments (Seydoux and Fire, 1994). Next, we compared the transcriptome of EMB PGCs to that of L1 PGCs to identify PGC transcripts whose abundance decline during embryogenesis. We identified 411 down-regulated transcripts, including 197 oocyte transcripts (Figure 2A, 2B and Table S5), consistent with turnover of many maternal mRNAs in PGCs after the 200-cell stage.Strikingly, the amplitude of this turnover was diminished in *nos-1(gv5)nos-2(RNAi)*mutants: the abundance of the 411 transcripts remained high overall during the transition from EMB PGCs to L1 PGCs in *nos-2(RNAi)nos-1(gv5)* embryos (Figure 2C).Furthermore, when comparing wild type and *nos-1(gv5)nos-2(RNAi)* EMB PGCs, we identified 182 differentially expressed transcripts (11 down- and 171 up-regulated), including 71 of oocyte transcripts that were more abundant in *nos-1(gv5)nos-2(RNAi)* EMB PGCs (Figure S2B). Together these findings suggest a defect in maternal mRNA turnover in *nos-1(gv5)nos-2(RNAi)* PGCs that is already detectable at the 200-cell stage and persist through embryogenesis. To test this hypothesis directly, we performed *in situ* hybridization experiments against three maternal mRNAs. In wild-type embryos, *mex-5*, C01G8.1 and Y51F10.2 are turned over rapidly in somatic lineages (~28-cell stage) and more slowly in the germ lineage (200-300-cell stage for *mex-5* and C01G8.1; bean-stage for Y51F10.2).We found that, in *nos-1(gv5)nos-2(ax3103)* embryos, turnover was not affected in somatic lineages, but was delayed in PGCs, with C01G8.1 persisting to the bean stage and *mex-5* and Y51F10.2 persisting to 1.5-fold stage (Figure 2D). We conclude that turnover of maternal mRNAs is compromised in *nos-1(gv5)nos-2(RNAi)* PGCs.

**Figure 2.**
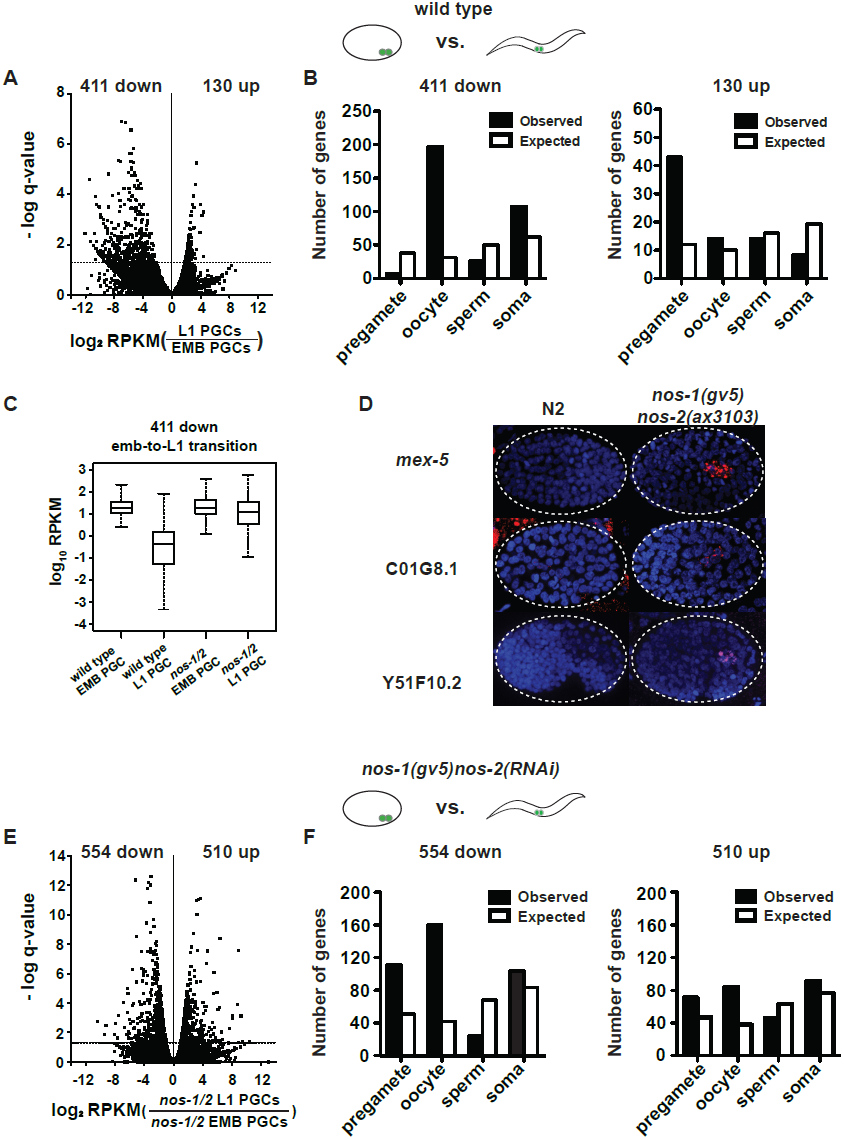
*nos-1nos-2* PGCs are defective in maternal mRNA turnover during embryogenesis. (A-B) Transcriptome comparison between PGCs isolated from wild-type embryos and wild-type L1 larvae. (A) Volcano plot showing log2 fold-change in transcript abundance for each gene. The numbers of genes whose expression were up- or down-regulated in L1 PGCs compared to embryonic PGCs are indicated. Dashed line marks the significance cutoff of q = 0.05 above which genes were counted as miss-expressed. (B) Bar graphs showing expected and observed number of genes (Y axis) in the different expression categories (X axis). The lists of expression categories used here are as in Figure 1B (Table S1). Oocyte genes are overrepresented among down-regulated genes and pre-gamete genes are overrepresented among up-regulated genes. (C) Box and whisker plot showing the expression levels (log10) of 411 genes that are down-regulated during embryogenesis in wild-type PGCs. Expression of these genes remains high on average in *nos-1(gv5)nos-2(RNAi)* PGCs. Each box extends from the 25th to the 75th percentile, with the median indicated by the horizontal line; whiskers extend from the 2.5th to the 97.5th percentiles. (D) Photomicrograph of embryos hybridized with single molecule fluorescence probes (red) against mex-5, C01G8.1 and Y51F10.2. Wild-type and nos-1(gv5)nos-2(ax3103) embryos were raised at 25°C and are compared here at the same stage (as determined by the number of DAPI-stained nuclei shown in blue). By the stages shown, all three transcripts have turned over in wild-type, but are still present (red signal) in PGCs in nos-1(gv5)nos-2(ax3103) embryos. At least ten embryos were examined per probe set in different genotypes shown. (E-F) Transcriptome comparison between PGCs isolated from *nos-1(gv5)nos-2(RNAi)*embryos and *nos-1(gv5)nos-2(RNAi)* L1 larvae. (E) Volcano plot showing log2 fold-change in transcript abundance for each gene. The numbers of genes whose expression were up- or down-regulated in L1 PGCs compared to embryonic PGCs are indicated. Dashed lines mark the significance cutoff of q = 0.05 above which genes were counted as miss-expressed. (F) Bar graphs showing expected and observed number of genes (Y axis) in the different expression categories (X axis).

### Transcription defects in PGCs lacking *nos-1* and *nos-2*

Comparison of the transcriptomes of EMB and L1 wild-type PGCs identified many transcripts whose abundance increase during embryogenesis, including large percentage of pre-gamete category (33%, 43/130, Figure 2A). In contrast, the same comparison in *nos-1(gv5)nos-2(RNAi)* PGCs identified upregulated genes in all categories, including 84 oogenic genes (Figure 2E, 2F and Table S5). These findings suggest that, by the L1 stage, *nos-1(gv5)nos-2(RNAi)* PGCs activate the transcription of many genes including oocyte genes, unlike wild-type L1 PGCs which primarily activate pre-gametic genes.

To explore this possibility further, we used ATAC-seq to identify regions of “open” chromatin that differ between wild-type and *nos-1(gv5)nos-2(RNAi)* L1 PGCs (Buenrostro et al., 2015). We first analyzed the ATAC-seq profile of 1430 genes that were upregulated in *nos-1(gv5)nos-2(RNAi)* L1 PGCs compared to wild-type (Figure 3A and Figure S1B-C). Metagene analysis identified a peak of increased chromatin accessibility at the transcriptional start site (Fig. 3A and B). The top 247 ATAC-seq+ genes (highest *nos-1/2*/WT peak ratio; Figure 3A, 3B and Table S2) were overexpressed in *nos-1(gv5)nos-2(RNAi)* L1 PGCs compared to wild-type (Figure 3C) and 106/247 were oogenic genes (Figure 3D). In contrast, 584 genes with decreased chromatin accessibility in *nos-1(gv5)nos-2(RNAi)* compared to wild-type had lower expression levels in *nos-1(gv5)nos-2(RNAi)* L1 PGCs compared to WT (Figure S3B, Table S2), and 187 corresponded to genes in the pre-gametic category (Figure S3C). These results suggest that PGCs lacking *nos-1nos-2* activate the transcription of oogenic genes that are silent in wild-type PGCs, and fail to fully activate the transcription of pre-gametic genes that are transcribed in wild-type PGCs.

**Figure 3.**
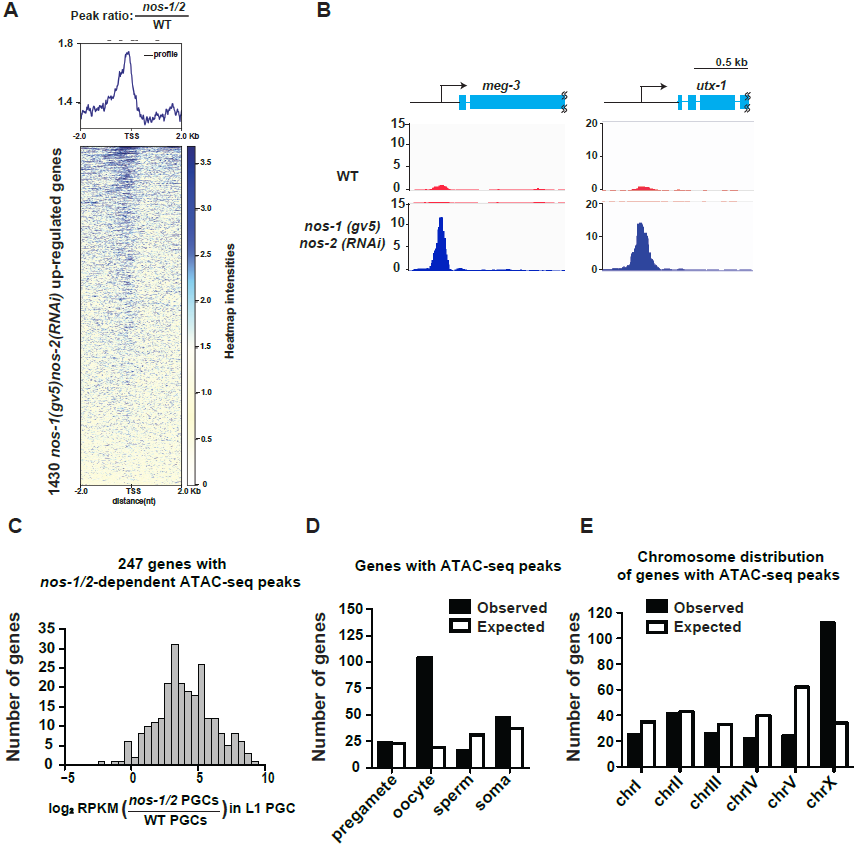
*nos-1nos-2* L1 PGCs activate the transcription of oocyte and X-linked genes. (A) Heat map showing accumulated ATAC-seq reads of 1430 genes (Y axis) that are upregulated genes in *nos-1(gv5)nos-2(RNAi)* L1 PGCs compared to wild-type (as determined by TruSeq, see Figure S1B and materials and methods). 4kb across transcription start site (TSS) are plotted on the heatmap (X axis). Darker color indicates accumulated reads (open chromatin) in *nos-1(gv5)nos-2(RNAi)*. See materials and methods for detailed description of the data analysis. See Table S5 for lists of genes with differential expression in PGCs. (B) Genome browser view of ATAC-seq profiles for *meg-3* and *utx-1*. Transcripts for both genes were significantly upregulated in *nos-1(gv5)nos-2(RNAi)* over wild-type L1 PGCs (See expression level in Table S4). The promoters of *meg-3* and *utx-1* are less accessible in wild-type compared to *nos-1(gv5)nos-2(RNAi)* PGCs. Y-axis shows normalized coverage read counts. (C) Histogram showing the distribution of log2 fold change in gene expression between *nos-1(gv5)nos-2(RNAi)* and wild-type L1 PGCs for 247 genes that acquired new ATAC-seq peaks in *nos-1(gv5)nos-2(RNAi)* PGCs (Table S2). Consistent with ATAC-seq peaks denoting open chromatin, most genes are expressed at higher levels in *nos-1(gv5)nos-2(RNAi)* PGCs compared to wild-type. (D) Bar graph showing expected and observed number of genes with *nos-1nos-2 -* dependent ATAC-seq peaks in the different expression categories. (E) Bar graph showing the chromosomal distribution of genes with *nos-1nos-2* -dependent ATAC-seq peaks.

Transcription of the X chromosome is silenced in all germ cells except in oocytes, which activate X-linked gene expression in preparation for embryogenesis (Kelly et al., 2002). As expected, we found that transcripts from X-linked genes are rare in wild-type L1 PGCs, with an average 4.7 FPKM per X-linked genes compared to 50.9 for autosomal genes. X-linked transcripts were more abundant in *nos-1(gv5)nos-2(RNAi)* L1 PGCs (9.6 FPKM for X-linked genes compared to 43.8 for autosomal genes) (Table S3), and strikingly 44% of the “open” genes with ATAC-seq peaks in *nos-1(gv5)nos-2(RNAi)* PGCs were X-linked (Figure 3E). We conclude that silencing of the X chromosome is defective in *nos-1(gv5) nos-2(RNAi)* PGCs.

### MES-2 and MES-4 activities are compromised in *nos-1nos-2* PGCs

Failure to silence X-linked genes has been reported for germ cells lacking the chromatin regulators *mes-2* and *mes-4* (Bender et al., 2006; Gaydos et al., 2012). To directly compare the effect of *nos* and *mes* activities in PGCs, we purified PGCs from L1 larvae derived from hermaphrodites where *mes-2* or *mes-4* was inactivated by RNAi (Methods). As expected, loss of *mes-2* and *mes-4* led to a significant upregulation of X-linked genes in L1 PGCs (Figure 4A-B, Figure S4A-B and Table S5 for lists of miss-regulated genes). To directly compare these changes to those observed in *nos-1(gv5)nos-2(RNAi)* PGCs, we compared, for each genotype, the log2 fold change over wild-type for X-linked genes and for autosomal oocyte genes. As expected, we observed a strong positive correlation between *mes-2* and *mes-4* in both gene categories (R=0.91 and R=0.76, X-linked and autosomal oogenic genes, respectively) (Figure 4C and 4D). We also observed a strong correlation between *mes-4(RNAi)* and *nos-1(gv5)nos-2(RNAi)*(R=0.75, Figure 4E) and *mes-2(RNAi)* and *nos-1(gv5)nos-2(RNAi)* (R=0.73, not shown) for X-linked genes. Interestingly, the correlations were weaker for autosomal oocyte genes (R=0.35, Figure 4F), which tended to be more over-expressed in *nos-1(gv5)nos-2(RNAi)* L1 PGCs. This finding is consistent with the notion that, while *nos-1nos-2* and *mes* PGCs share a defect in X-linked silencing, *nos-1nos-2* PGCs also have an additional defect in maternal mRNA turn over.

**Figure 4.**
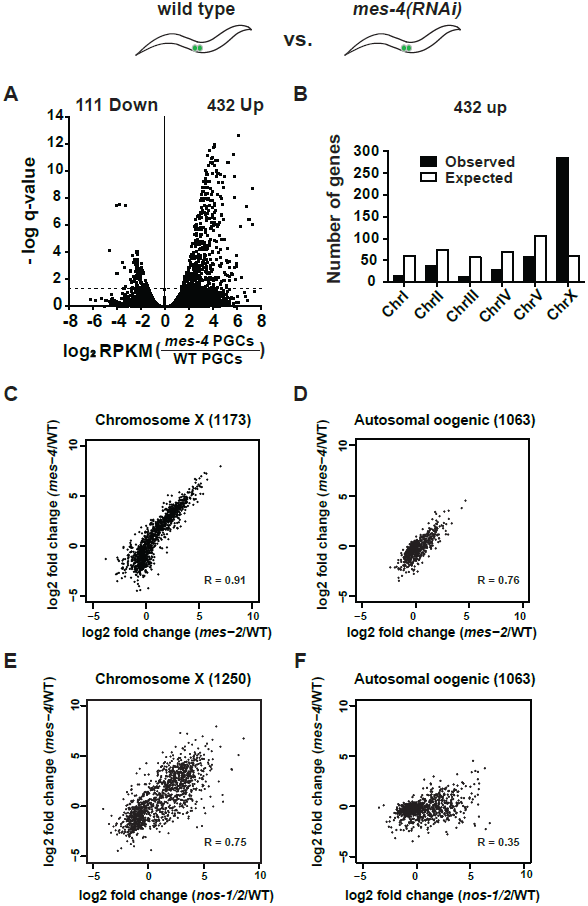
*nos-1nos-2* PGCs share a defect in X chromosome silencing with *mes-4* PGCs. (A-B) Transcriptome comparison between PGCs isolated from wild-type and *mes-4(RNAi)* L1 larvae. See Fig. S4 for comparison between wild-type and *mes-2(RNAi)*. (A) Volcano plot showing log2 fold-change in transcript abundance for each gene. The numbers of genes whose expression were up- or down-regulated in *mes-4(RNAi)* PGCs are indicated. Dashed lines mark the significance cutoff of q = 0.05 above which genes were counted as miss-expressed. (B) Bar graph showing chromosomal distribution of *mes-4(RNAi)* up-regulated genes. (C-D) XY scatter plots showing correlation of the fold change in gene expression between *mes-2(RNAi)* (X-axis) and *mes-4(RNAi)*(Y-axis) PGCs compared to wild-type for X-linked genes and autosomal oogenic genes. Pearson‘s correlation values are indicated. (E-F) XY scatter plots showing correlation of the fold change in gene expression between *nos-1(gv5)nos-2(RNAi)* (X-axis) and *mes-4(RNAi)* (Y-axis) PGCs compared to wild-type for X-linked genes and autosomal oogenic genes. Pearson‘s correlation values are indicated.

MES-2, 3, 4 and 6 proteins are maternally-inherited and are maintained in PGCs throughout embryogenesis (Fong et al., 2002; Holdeman et al., 1998; Korf et al., 1998; Strome, 2005). We observed no significant changes in *mes* transcripts in *nos-1(gv5)nos-2(RNAi)* PGCs compared to wild-type (Table S4). Direct examination of MES-2, MES-3 and MES-4 proteins confirmed that their expression patterns were unchanged in *nos-1(gv5)nos-2(ax3103)* or *nos-1(gv5)nos-2(RNAi))* embryos (Figure S4C). Together, these results suggest that *nos-1* and *nos-2* do not affect MES expression directly despite being required for MES-dependent silencing.

### Loss of *lin-15B, lin-35/RB* and *dpl-1/DP* suppresses *nos-1nos-2* sterility

MES-dependent silencing in somatic cells and adult germlines is antagonized by members of the synMuvB class of transcriptional regulators (Petrella et al., 2011; Tabuchi et al., 2013). To test whether synMuvB activity contributes to the *nos-1nos-2* PGC phenotype, we tested whether inactivation of synMuvB genes could reduce the sterility of *nos-1nos-2* animals using combinations of RNAi and mutants (Figure S5) and verified positives by analyzing the sterility of triple mutant combinations (Figure 5). We found that loss-of-function mutations in *lin-15B, lin-35* and *dpl-1* reduced the sterility of *nos-1(gv5)nos-2(ax3103)* from >70% to <30%. (Figure 5A). The most dramatic reduction was seen with *lin-15B(n744)*, which reduced *nos-1(gv5)nos-2(ax3103)* sterility to 3.4% (Figure 5A). *lin-15B* is a THAP domain DNA binding protein that has been implicated with the DRM class of transcriptional regulators, including *lin-35* and *dpl-1*, in the silencing of germline genes in somatic cells (Araya et al., 2014; Petrella et al., 2011; Wu et al., 2012). Other DRM components (*efl-1, lin-37, lin-9, lin-52, lin-54*), however, did not suppress *nos-1(gv5)nos-2(ax3103)* sterility (Figure S5A).

**Figure 5.**
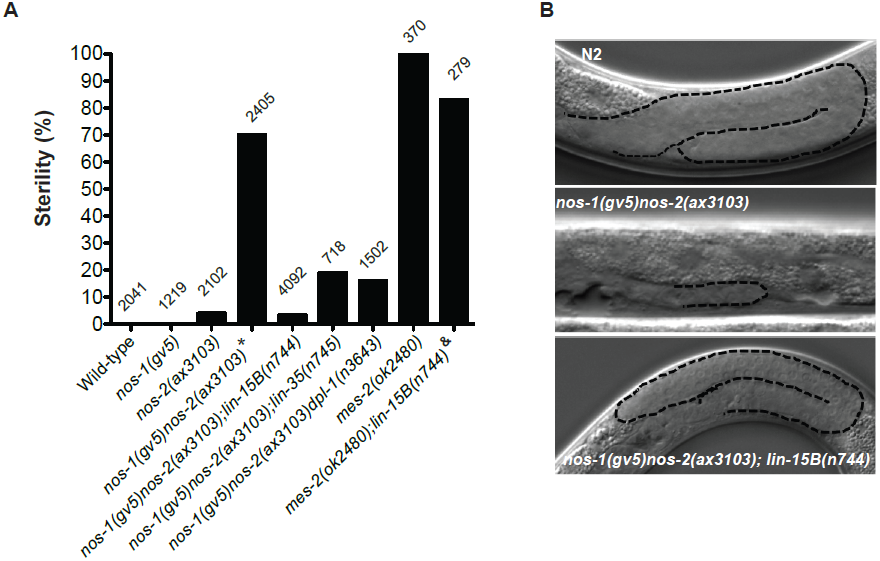
Suppression of *nos-1nos-2* sterility by *synMuvB* mutants. (A) Bar graph showing sterility of progenies from listed genotypes. *lin-15B(n744)* and *lin-35(n745)* are null alleles (Ferguson and Horvitz (1989), Lu and Horvitz (1998), Petrella et al.(2011)). *dpl-1(n3643)* is a loss of function allele (Ceol and Horvitz (2001), Petrella et al.(2011)). *mes-2(ok2480)* is a deletion allele that causes 100% maternal-effect sterility (Consortium et al., 2012). Number of hermaphrodites scored is written above indicated genotypes. ^*$*^ *nos-1(gv5)nos-2(ax3103)* hermaphrodites produce 70% sterile progenies at 20°C and 96% sterile progeny at 25ଌ with severely atrophied germlines (Figure 5B). *nos-1(gv5)nos-2(ax3103)*; *lin-15(n744)* hermaphrodites produce 96.6% fertile progenies at 20ଌ, and arrest as larvae at 26ଌ as is true of *lin-15(n744)* animals. *& mes-2(ok2480); lin-15(n744)* hermaphrodites cannot be maintained as a selfing population (Figure S5). (B) Nomarski Images of germlines (stippled) in L4 hermaphrodites of the indicated genotypes. Worms were staged according to vulval development.

Since PGCs lacking *mes* and *nos-1nos-2* shared the same defect in X chromosome silencing (Figure 4E), we tested whether loss of *lin-15B* could also suppress *mes-2* maternal effect sterility. Hermaphrodites derived from *mes-2(ok2480)* mothers are 100% sterile (Figure 5A and S5B). We found that *lin-15B(n744)* suppressed *mes-2(ok2480)* sterility weakly and only for one generation. Animals derived from *mes-2(ok2480); lin-15B(n744)* mothers were 83% sterile in the first generation and 98% sterile in the second generation and could not be maintained as a selfing population (Figure 5A and S5B). In contrast, *nos-1(gv5)nos-2(ax3103); lin-15B(n744)* triple mutant animals were almost fully fertile (96.6% fertile, Figure 5A) and could be maintained as a selfing population for >10 generations. We conclude that inactivation of *lin-15B* bypasses the requirement for *nos-1nos-2* activity, but not for *mes* activity.

### Maternal LIN-15B is inherited by all embryonic blastomeres and downregulated specifically in PGCs

A LIN-15B::GFP transgene was reported to be broadly expressed (Sarov et al., 2012). To examine the expression of endogenous LIN-15B, we used a polyclonal antibody generated against LIN-15B protein (modencode project, personal communication with Dr. Susan Strome). We confirmed the specificity of this antibody by staining *lin-15B(n744)* mutant, which showed no nuclear staining (Figure S6A). We first detected LIN-15B expression in the germline in the L4 stage in nuclei near the end of the pachytene region where germ cells initiate oogenesis (Figure 6A). Nuclear LIN-15B was present in all oocytes and inherited by all embryonic blastomeres, including the germline P blastomeres (Figure 6B, S6A). LIN-15B remained present at high levels in all somatic nuclei throughout embryogenesis. In contrast, in the germ lineage, LIN-15B levels decreased sharply during the division of the germline founder cell P4 that generates the two PGCs (Figure 6B Left panels). LIN-15B expression remained at background levels in PGCs throughout embryogenesis.

**Figure 6.**
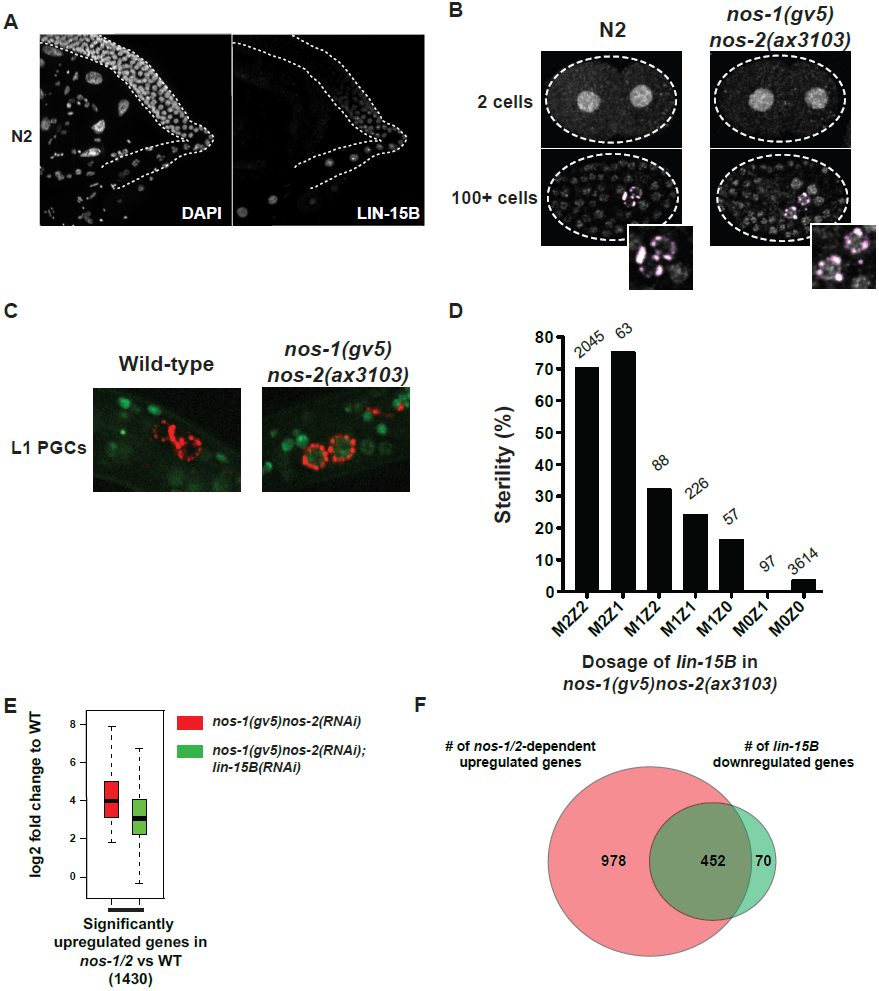
LIN-15B is inherited maternally and is downregulated in PGCs in a *nos-1nos-2* dependent manner. (A) Photomicrograph of dissected wild-type gonad stained for anti-LIN-15B antibody and DAPI for DNA. LIN-15B protein is detected at the end of the pachytene region and in all oocyte nuclei. (B) Photomicrographs of fixed wild-type and *nos-1(gv5)nos-2(ax3103)* embryos stained with anti-LIN-15B and K76 (anti-PGL-1, red) antibodies. The anti-LIN-15B polyclonal serum cross-reacts with perinuclear germ granules (pink color, see Methods). 45/60 PGCs were positive for LIN-15B in *nos-1(gv5)nos-2(ax3103)* embryos compared to 0/34 in wild-type. (C) Photomicrographs of newly hatched gonads from wild-type and *nos-1(gv5)nos-2(ax3103)* L1 larvae with a paternal copy of the *lin-15B* transcriptional reporter (green). 12/16 *nos-1(gv5)nos-2(ax3103)* PGC doublets were positive for GFP compared to 0/28 in wild-type. (D) Bar graph showing the sterility of *nos-1(gv5)nos-2(ax3103)* hermaphrodites with different dosages of maternal and zygotic *lin-15B*. Mating schemes are shown in Figure S7. Number of hermaphrodites scored is written above each genotype. (E) Box and Whisker plot showing log2 fold change compared to wild-type of 1430 genes that are upregulated in *nos-1(gv5)nos-2(RNAi)* L1 PGCs. Each box extends from the 25th to the 75th percentile, with the median indicated by the horizontal line; whiskers extend from the 2.5th to the 97.5th percentiles. The upregulation is reduced in *nos-1(gv5)nos-2(RNAi);lin-15B(RNAi)* PGCs. (F) Venn diagram showing overlap between 1430 genes upregulated in *nos-1(gv5)nos-2(RNAi)* compared to wild-type L1 PGCs (red) and downregulated genes in *nos-1(gv5) nos-2(RNAi*); *lin-15B(RNAi)* compared to *nos-1(gv5)nos-2(RNAi)* L1 PGCs (522 genes, green).

### Downregulation of maternal LIN-15B in PGCs requires *nos-1 nos-2* activity

*lin-15B* transcripts were modestly elevated in *nos-1(gv5)nos-2(RNAi)* EMB PGCs compared to wild-type EMB PGCs, suggesting that *lin-15B* may be one of the maternal RNAs that requires Nanos activity for rapid turnover in PGCs (Table S4). *lin-15B* transcripts rose significantly by ~2-fold when comparing EMB versus L1 stage *nos-1(gv5)nos-2(RNAi)* PGCs. This increase was not observed in wild-type PGCs, suggesting that *lin-15B* is also inappropriately transcribed in *nos-1(gv5)nos-2(RNAi)* L1 PGCs. Unfortunately, we were not able to confirm these RNA-seq observations by *in situ* hybridization due to the low abundance of *lin-15B* RNA and its presence in all somatic cells.

To determine whether LIN-15B protein expression is under the control of Nanos activity, we stained embryos derived from *nos-1(gv5)nos-2(ax3103)* hermaphrodites with the anti-LIN-15B antibody. We found that, in contrast to wild-type, *nos-1(gv5)nos-2(ax3103)* embryos maintained high LIN-15B levels in embryonic PGCs (Figure 6B Right panels).*nos-1(gv5)nos-2(ax3103)* embryos could miss-regulate LIN-15B by delaying the turnover of maternal LIN-15B or by activating premature zygotic transcription of the *lin-15B* locus. To distinguish between these possibilities, we created a *lin-15B* transcriptional reporter by inserting a GFP::H2B fusion at the 5‘ end of *lin-15B* locus in an operon configuration to preserve endogenous *lin-15B* expression (Figure S6B and Table S6). We crossed *nos-1(gv5)* males carrying the *lin-15B* transcriptional reporter to wild-type or *nos-1(gv5)nos-2(ax3103)* hermaphrodites and examined crossed progenies for GFP expression. In both cases, we observed strong GFP expression in somatic cells, but no expression in PGCs during embryogenesis (data not shown). In wild-type animals, we first observed zygotic expression of the *lin-15B* transcriptional reporter in the germline of L4 stage animals (Figure S6C), in germ cells that have initiated oogenesis. In contrast, in animals derived from *nos-1(gv5)nos-2(ax3103)* mothers, zygotic expression of the *lin-15B* transcriptional reporter could be detected as early as the L1 stage in PGCs and their descendants (Figure 6C). This expression was maintained until the L2 stage when *nos-1nos-2* PGC descendants undergo cell death. We conclude that *nos-1nos-2* activity is required both to promote the turnover of maternal LIN-15B in EMB PGCs and to prevent premature zygotic transcription of *lin-15B* in L1 PGCs.

### Maternal *lin-15B* is responsible for *nos-1 nos-*2 sterility

To determine whether miss-regulation of maternal or zygotic LIN-15B is responsible for *nos-1nos-2* sterility, we compared the sterility of *nos-1(gv5)nos-2(ax3103)* animals that lack either maternal or zygotic *lin-15B* (Figure 6D and Figure S7). We found that loss of maternal *lin-15B* was sufficient to fully suppress *nos-1(gv5)nos-2(ax3103)* sterility, even in the presence of one zygotic copy of *lin-15B* (Figure 6D). The penetrance of the suppression was dependent on the dosage of maternal *lin-15B*. *nos-1(gv5)nos-2(ax3103)* animals with only one copy of maternal *lin-15B* were only 32% sterile compared to 70% sterility for animals with two copies of maternal *lin-15B* and 0% with animals with zero copies of maternal *lin-15B* (Figure 6D). Interestingly animals with only one copy of maternal LIN-15B appeared sensitive to the zygotic dosage of *lin-15B* (Figure 6D, compare the sterility M1Z2, M1Z1 and M1Z0). We conclude that maternal *lin-15B* is primarily responsible for the sterility of *nos-1nos-2* animals, although zygotic LIN-15B activity may also contribute.

### Loss of *lin-15B* activity mitigates gene expression changes in *nos-1 nos-2* PGCs

LIN-15B is a transcription factor with many targets in somatic cells but no known function in the germline (Niu et al., 2011). To determine the effect of ectopic LIN-15B on the transcriptome of *nos-1(gv5)nos-2(RNAi)* PGCs, we profiled *nos-1(gv5)nos-2(RNAi); lin-15B(RNAi)* PGCs and compared the log2 fold change of transcripts in *nos-1(gv5)nos-2(RNAi); lin-15B(RNAi)* PGCs and *nos-1(gv5)nos-2(RNAi)* PGCs to wild-type. We found that loss of *lin-15B* reduced gene miss-expression in *nos-1(gv5)nos-2(RNAi)* PGCs (Figure 6E). Of the 1430 upregulated genes in *nos-1(gv5)nos-2(RNAi)* PGCs, 31% (452) had significantly lower expression levels in *nos-1(gv5)nos-2(RNAi); lin-15B(RNAi)* PGCs (Figure 6F). Both upregulated and down-regulated gene categories were rescued, as well as X-linked and oogenic genes (Figure S6D). These data indicate that ectopic *lin-15B* activity is responsible for a significant number of miss-expressed genes in *nos-1(gv5)nos-2(RNAi)* PGCs.

We compared the lists of genes upregulated in *nos-1(gv5)nos-2(RNAi)* and *mes-4(RNAi)* PGCs and down-regulated in *nos-1(gv5)nos-2(RNAi); lin-15B(RNAi)* PGCs (Table S5). We identified 88 shared genes and 70 of them are X-linked genes, including *utx-1*. *utx-1* encodes a X-linked histone demethylase specific for the H3K27me3 mark generated by *mes-2* (Agger et al., 2007; Seelk et al., 2016). Like other X-linked genes, *utx-1* transcripts are rare in wild-type PGCs (FPKM<0.2)(Table S4) and are overexpressed 9.1-fold in *mes-2(RNAi)* PGCs. In *nos-1(gv5) nos-2(RNAi)* L1 PGCs, the *utx-1* locus acquires a new ATAC-seq peak (Figure 3B) and *utx-1* transcripts are overexpressed 160-fold. This overexpression was reduced significantly by 2.4-fold in *nos-1(gv5)nos-2(RNAi); lin-15B(RNAi)* PGCs. These observations suggest that *utx-1* may function downstream of *lin-15B* to further antagonizes MES activity as X-linked genes become desilenced. If so, loss of *utx-1* should alleviate *nos-1nos-2* sterility. Consistent with this prediction, we found that reduction of *utx-1* activity by RNAi partially suppressed *nos-1(gv5)nos-2(ax3103)* sterility (Figure S5A). Suppression by *utx-1* was not as extensive as that observed with *lin-15B*, suggesting that *utx-1* is not the only gene activated in *nos-1(gv5)nos-2(ax3103)* PGCs that leads to sterility. We conclude that activation of *utx-1* by ectopic *lin-15B* is at least partially responsible for the loss of *mes* activity and sterility observed in *nos-1nos-2* animals.

## Discussion

In this study, we have examined the transcriptome of PGCs lacking Nanos function in *C. elegans*. We have found that *nos-1(gv5)nos-2(RNAi)* PGCs activate 100s of genes normally expressed in oocytes and somatic cells. Our observations suggest that Nanos activity is required to erase a maternally-inherited somatic program. Importantly, Nanos activity promotes the turnover of LIN-15B, a maternally-inherited transcription factor known to antagonize PRC2 activity in somatic cells. Down-regulation of LIN-15B frees PRC2/MES-4 to silence oocyte and somatic genes and activate germline genes in PGCs.

### Nanos activity is required for the timely turnover of maternal mRNAs in PGCs

During oogenesis, oocytes stockpile mRNAs and proteins in preparation for embryogenesis. These include mRNAs and proteins with housekeeping functions as well as factors required to specify somatic and germ cell fates. During the maternal-to-zygotic transition, these maternal products are turned over to make way for zygotic factors. We have found that Nanos activity is required for the timely turnover of maternal mRNAs in PGCs. RNA-seq analyses comparing embryonic and first stage larval PGCs identified 411 maternal mRNAs whose abundance decrease sharply during embryogenesis in wild-type PGCs, but not in *nos-1(gv5)nos-2(RNAi)* PGCs (Figure 2E). We also found that maternal LIN-15B protein levels decline rapidly at the time of gastrulation in wild-type PGCs, but not in *nos-1(gv5)nos-2(ax3103)* PGCs (Figure 6B). Nanos is thought to silence mRNAs by interacting with the sequence-specific RNA-binding protein Pumilio and with the CCR4-NOT deadenylase complex which interferes with translation and can also destabilize RNAs. (Lai et al., 2012; Suzuki et al., 2012; Swartz et al., 2014; Wharton et al., 1998). In the *C. elegans* genome, there are eight genes related to *Drosophila pumilio*. Depletion of five of these (*fbf-1, fbf-2, puf-6, puf-7 and puf-8)* phenocopies the *nos-1nos-2* PGC phenotypes, including failure to incorporate in the somatic gonad, premature proliferation, and eventually cell death (Subramaniam and Seydoux, 1999). These observations suggest that NOS-1 and NOS-2 function with Pumilio-like proteins to target specific maternal RNAs for degradation. Paradoxically, in sea urchins, Nanos targets the mRNA coding for the CNOT6 deadenylase for degradation in PGCs, which indirectly stabilizes other maternal mRNAs (Swartz et al., 2014). In that system, Nanos was also found to silence eEF1A expression, leading to a transient period of translational quiescence in PGCs (Oulhen et al., 2017).One possibility is that, at the earliest stages of the maternal-to-zygotic transition in PGCs, Nanos generally silences maternal mRNA translation and targets specific mRNAs for degradation while stabilizing others. In combination, these effects could lead to loss of somatic mRNAs and proteins (e.g. LIN-15B) and maintenance of germline mRNAs (e.g. MES) whose translation could be reactivated at a later time. In *C. elegans*, the redundant *nanos* homologs *nos-1* and *nos-2* are expressed sequentially in PGCs during the maternal-to-zygotic transition and may have overlapping yet distinct effects on mRNAs stability and translation. Genetic analyses already have suggested that *nos-1* and *nos-2*have both unique and shared functions (Kapelle and Reinke, 2011; Mainpal et al., 2015). It will be important to determine whether *nos-1* and *nos-2* are both required to keep LIN-15B levels low throughout embryogenesis, and whether they act directly on the *lin-15B* RNA or indirectly, by silencing other factors required for LIN-15B protein translation and/or stability.

### In PGCs lacking *nos-1* and *nos-2*, maternal LIN-15B interferes with MES-dependent silencing of oocyte and somatic genes

Several lines of evidence indicate that *nos1nos-2* sterility is caused by a failure to turn over maternally-inherited LIN-15B in embryonic PGCs. First, loss of one maternal copy of the *lin-15B* locus is sufficient to partially suppress *nos-1(gv5)nos-2(ax3103)* sterility and loss of both maternal copies maximally suppresses *nos-1(gv5)nos-2(ax3103)* sterility even in the presence of a zygotic copy of *lin-15B* (Figure 6D). These genetic results demonstrate that maternal *lin-15B* is required for *nos-1(gv5)nos-2(ax3103)* sterility, and suggest that abnormal perdurance of LIN-15B in PGCs interferes with their reprogramming to become pre-gametic germ cells. Consistent with the genetic findings, we have found that *nos-1(gv5)nos-2(RNAi)* PGCs activate by the L1 stage the transcription of 100s of somatic and oocyte genes and this ectopic expression is reduced in PGCs also lacking *lin-15B* activity (Figures 1 and 6). How does ectopic LIN-15B activate oocyte and somatic gene expression in *nos-1nos-2* PGCs? LIN-15B activity antagonizes MES-dependent repression of somatic genes and activation of germline genes in somatic cells (Petrella et al., 2011; Wang et al., 2005). Consistent with LIN-15B playing a similar role in PGCs, *nos-1(gv5)nos-2(RNAi)* PGCs activate the transcription of many of the same genes activated in PGCs lacking *mes* activity. The strongest correlation is seen for genes on the X chromosome (Figure 4E), a well-documented focus of MES transcriptional repression (Bender et al., 2006; Garvin et al., 1998; Gaydos et al., 2012). Interestingly, the *lin-15B* locus itself is on the X chromosome and is ectopically transcribed in *nos-1(gv5)nos-2(RNAi)* PGCs at hatching. These observations raise the possibility that maternal LIN-15B potentiates zygotic *lin-15B* expression as MES-dependent silencing of the X-chromosome becomes compromised. How does maternal LIN-15B initially opposes MES activity is not known, but another X-linked gene and potential LIN-15B target is *utx-1*, a de-methylase that removes the silencing mark deposited by the PRC2 complex. Upregulation of *utx-1* was shown recently to promote reprogramming of adult germline stem cells into neurons (Seelk et al., 2016). *utx-1* is up-regulated in a *lin-15B*-dependent manner in *nos-1(gv5)nos-2(RNAi)* PGCs, and RNAi of *utx-1* partially suppresses *nos-1(gv5)nos-2(ax3103)* sterility (Figure S5A). Suppression by loss of *utx-1* is weaker than that observed when inactivating *lin-15B*, suggesting that *utx-1* is not the only *lin-15B* target that opposes PRC2. We have found that loss of two other synMuvB genes *lin-35/Rb* and *dpl-1* also suppresses *nos-1(gv5)nos-2(ax3103)* sterility (Figure 5A), albeit again less stringently than loss of *lin-15B*. It will be interesting to determine whether these genes function with, or in parallel to, LIN-15B to oppose PRC2 activity in PGCs.

Inhibition of LIN-15B by Nanos is unlikely to be the only mechanism that promotes PCR-2 function in PGCs. XND-1 is a chromatin-associated protein that is expressed in PGCs throughout embryogenesis. XND-1 is required redundantly with NOS-2 to maintain low levels of active histone marks in PGCs (Mainpal et al., 2015). An exciting possibility is that XND-1 is a chromatin factor that promotes/maintains PRC2 activity in PGCs, in parallel to NOS-2.

### An ancient regulatory switch balances somatic and germline fates throughout the germline cycle

Competition between synMuvB and PRC2 activities has already been implicated in balancing somatic and germline gene expression during larval development in somatic lineages and in the adult germline (Petrella et al., 2011; Tabuchi et al., 2013). Our findings demonstrate that such a competition also occurs in PGCs, where Nanos biases the competition in favor of PRC2 by lowering maternal LIN-15B levels (Figure 7A). We propose that the ratio of synMuvB and PRC2 activity changes at two key developmental stages during the germline cycle (Figure 7B). First, during oogenesis, an unknown activity promotes the transcriptional activation of LIN-15B, which allows the demethylase UTX-1 and other LIN-15B targets to begin erasing PRC2 marks. Erasure of PRC2 marks activates the transcription of X-linked genes and other somatic genes in oocytes in preparation for embryogenesis. This oogenic/maternal program is inherited by all embryonic blastomeres. In somatic lineages, which activate transcription first, maternally-inherited LIN-15B continues to oppose PRC2 activity, which permits zygotic activation of the *lin-15B* and *utx-1* loci and eventual complete erasure of the PRC2 program. In the nascent germline, transcription is kept off until gastrulation when Nanos expression is activated in PGCs by unknown mechanisms that both promote the translation of maternal *nos-2* RNA and later the zygotic transcription of *nos-1*. Nanos activity in PGCs promotes the turnover of maternal LIN-15B, freeing PRC2 to re-establish silencing of somatic and X-linked genes, including the *lin-15B* and *utx-1* loci, until the next round of oogenesis.

**Figure 7.**
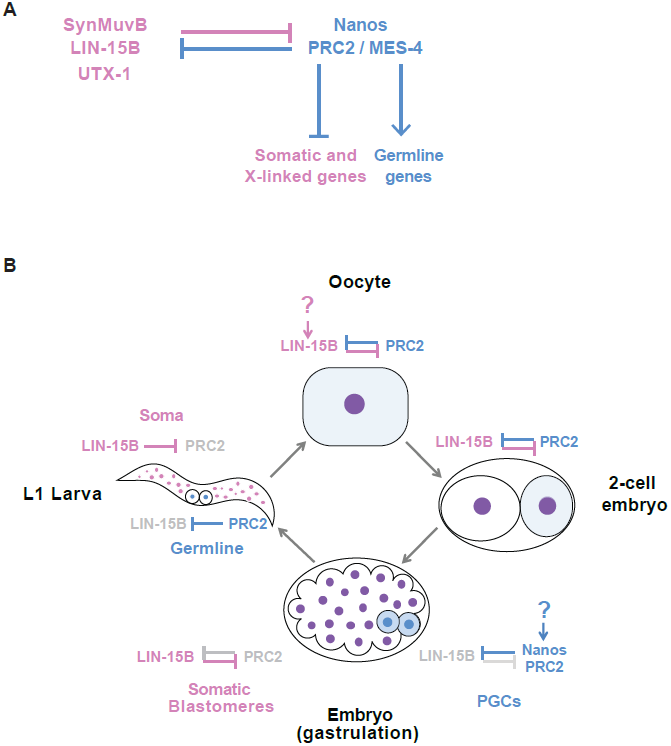
Mutual antagonism model. (A) Working model: A cross-regulatory loop balances activities that promote somatic (pink) and germline (blue) gene expression. LIN-15B, and other factors including UTX-1, opposes PRC2(Petrella et al., 2011). PRC2 silences somatic genes and X-linked genes (including *lin-15B*) and activates germline genes (with the help of MES-4) (Gaydos et al., 2012). Nanos post-transcriptionally down-regulates LIN-15B (this study). (B) Working model: Mutual antagonism between LIN-15B and PRC2 balances somatic and germline fates during development. In oocytes, LIN-15B transcription is activated by an unknown mechanism, leading to co-expression of LIN-15B and PRC2 in oocyte nuclei (purple). Competition between LIN-15B and PRC2 begins to erase PRC2 silencing marks, allowing the activation of somatic and X-linked genes in oocytes. In embryos, maternal LIN-15B and PRC2 are co-inherited (purple) by all nuclei. Somatic blastomeres activate zygotic transcription early when maternal LIN-15B levels are still high, causing the complete erasure of PRC2 marks and zygotic activation of somatic and X-linked genes,including *lin-15B*. In germline blastomeres, the onset of zygotic transcription is delayed until gastrulation by maternal proteins (light blue) that segregate with the nascent germline and also stabilize and promote the translation of maternal RNAs such as *nos*-2 (Seydoux and Braun, 2006; Tenenhaus et al., 2001). Nanos activity promotes the turnover of maternal LIN-15B, leaving PRC2 (blue nuclei) free to silence somatic and X-linked genes, including *lin-15B*. At hatching, somatic cells have high LIN-15B activity and PGCs have high PRC2 activity.

What prevents expression of Nanos in somatic cells? Interestingly, evidence in Drosophila and mammals suggest that Nanos is among the germline genes inhibited by synMuvB activity in somatic cells. Loss of the dREAM complex component *lethal (3)malignant brain tumor [ l(3)mbt]* leads to tumorous growth in Drosophila imaginal disks and ectopic expression of germline genes, including *nanos* (Janic et al., 2010). Similarly, loss of the synMuv B class transcription factor retinoblastoma protein (RB) leads to activation of *nanos* transcription in mammalian tissue culture cells and in *Drosophila* wings (Miles and Dyson, 2014; Miles et al., 2014). A complex regulatory feedback loop has also been reported between the LSD1 demethylase and the Nanos partner Pumilio in Drosophila and human bladder carcinoma cells (Miles et al., 2015). Taken together, these observations suggest that mutual antagonism between transcriptional regulators and the Nanos/Pumilio families of RNA binding proteins may be part of an ancient cross-regulatory loop that balances somatic and germline gene expression during development by controlling PRC2 activity. Key questions for the future will be to understand how the switch is flipped from germline-to-soma in oocytes (what downregulates Nanos expression in oocytes and activate LIN-15B expression?), from soma-to-germline in PGCs (what activates Nanos expression and how does Nanos down-regulate LIN-15B?), and how the switch becomes deregulated in malignancies.

## Acknowledgments

We thank the UC riverside IIGB Genomics Core facility, the JHU GRCF sequencing facility, and flow cytometry Core Facilities at Johns Hopkins Sidney Kimmel Comprehensive Cancer Center and the Johns Hopkins Bloomberg school of public health for expert support. We thank members in Seydoux lab and Baltimore worm club for discussions. We thank Dr. Sijung Yun at Yotta Biomed, LLC for his assistance and discussions in data analysis. We thank Dr. Julie Ahringer and Dr. Yan Dong for their assistance in ATAC-seq procedure. We also thank Dr. Susan Strome for sharing results of gene category analysis associated with Gaydos et al., 2012. This work was supported by National Institutes of Health (NIH) [grant number R01HD37047]. G.S. is an investigator of the Howard Hughes Medical Institute. Chih-Yung S Lee is a Damon Runyon Fellow supported by the Damon Runyon Cancer Research Foundation (DRG-2417-13).

## Materials and methods

### Worm handling, RNAi, sterility counts

*C. elegans* was cultured according to standard methods (Brenner, 1974).

RNAi knock-down experiments were performed by feeding on HT115 bacteria (Timmons and Fire, 1998). Feeding constructs were obtained from Ahringer or OpenBiosystem libraries or PCR fragments cloned into pL4440. The empty pL4440 vector was used as negative control. Bacteria were grown at 37ଌ in LB + ampicillin (100 μg/mL) media for 5-6 hr, induced with 5 mM IPTG for 30 min, plated on NNGM (nematode nutritional growth media) + ampicillin (100 μg/mL) + IPTG (1 mM) plates, and grown overnight at room temperature. Embryos isolated by bleaching gravid hermaphrodites, or synchronized L1s hatched in M9, were put onto RNAi plates. For sterility counts, the progeny of at least six gravid adult hermaphrodites were tested. Adult progenies were scored for empty uteri (‘white sterile’ phenotype) on a dissecting microscope. For all Immunostaining and smFISH experiments shown in Figure 2D, 6A, 6B, S4C and S6A, worms were grown at 25°C. For live embryo imaging and synMuvB related experiments shown in Figure 5, Figure 6C, 6D and Figure S4C, worms were grown at 20°C.

To verify the efficiency of RNAi treatments used to create sequencing libraries, we scored animals exposed to the same RNAi feeding conditions for maternal-effect sterility. For *nos-1(gv5)* strain on *nos-2* RNAi, sterility was 81%±10% at 20°C; and 86%±6% at 25°C; mes-2*(RNAi)* maternal effect sterility was 51%±1.4% and *mes-4(RNAi)* maternal effect sterility was 95.5%±3.5%. To test the efficiency of the double RNAi treatment for *nos-1(gv5)nos-2(RNAi); lin15B(RNAi)* RNA-seq libraries, we performed two additional controls. First we exposed a *nos-2::FLAG* strain (Paix et al., 2014) to the same RNAi feeding conditions and stained the embryos with anti::FLAG antibody to confirm knock down of *nos-2* (4/15 embryos showed weak staining, compared to 15/15 embryos with strong staining in the untreated controls). Second, we exposed *a lin-15B::GFP* strain (Paix et al.,2014) to the same double RNAi feeding conditions and observed no GFP expression in embryos. *nos-1(gv5)nos-2(RNAi); lin15B(RNAi)* animals gave 34%±19% sterile progeny.

## Generation of *nos-2* null allele by CRISPR-mediated genome editing

See Supplemental Tables S6 (strain table) and Table S7 (Oligo table) for lists of strains and CRISPR reagents. The *nos-2(ok230)* allele removes the *nos-2* coding region and a flanking exon in the essential gene *him-14*, resulting in embryonic lethality. To create a *nos-2* null allele that does not affect *him-14* function, we deleted the *nos-2* open reading frame using CRISPR/Cas9-mediated genome editing (Paix et al., 2015). Consistent with previous reports (Mainpal et al., 2015; Subramaniam and Seydoux, 1999), *nos-2(ax3103)* animals are viable and fertile and *nos-1(gv5) nos-2(ax3103)* double mutants are maternal effect sterile (Figure 5A).

### Immunostaining

Adult worms were placed on 3-wells painted slides in M9 solution (Erie Scientific co.) and squashed under a coverslip to extrude embryos. Slides were frozen by laying on pre-chilled aluminum blocks for >10 min. Embryos were permeabilized by freeze-cracking (removal of coverslips from slides) followed by incubation in methanol at −20°C; for 15 min, and then in pre-chilled acetone at −20°C; for 10 min. Slides were blocked twice in PBS-Tween (0.1%)-BSA (0.1%) for 15 min at room temperature, and incubated with 75 μml primary antibody overnight at 40°C in a humidity chamber. Antibody dilutions (in PBST/BSA): Rabbit α-LIN-15B 1:20,000 (SDQ3183, gift from Dr. Susan Strome), Rabbit α-MES-4 1:400 (Gift from Dr. Susan Strome,), K76 (1:10, DSHB), Rat α-OLLAS-L2 (1:200, Novus Biological Littleton, CO), Rat α-OLLAS 1:80 (Gift from Dr. Jeremy Nathans), mouse anti-FLAG M2 1:500 (Sigma F3165). Secondary antibodies (Molecular Probes/Thermo Fisher Sci.) were applied for 1~2 hr at room temperature. MES-3 was tagged with the OLLAS epitope at the C-terminus using CRISPR genome editing (Paix et al., 2015).

### Confocal microscopy

Fluorescence microscopy was performed using a Zeiss Axio Imager with a Yokogawa spinning-disc confocal scanner. Images were taken and stored using Slidebook v6.0 software (Intelligent Imaging Innovations) using a 40x or 63x objective. Embryos were staged by DAPI-stained nuclei in optical Z-sections and multiple Z-sections were taken to include germ cells marked by anti-PGL-1 (K76) staining. For images of embryonic PGCs, a single Z-section was extracted at a plane with the widest area of DAPI staining for nuclear signal of LIN-15B, MES-3, and MES-4. For MES-2-GFP, the Z-section was determined based on widest area of GFP signal. Equally normalized images were first taken by Slidebook v6.0, and contrasts of images were equally adjusted between control and experimental sets using Image J.

### Germ cell isolation and sorting

RNAi treatments for sorting experiments were done by seeding synchronized L1 (hatched from embryos incubated in M9 overnight) onto RNAi plates and growing them to gravid adults. Additional RNAi or control bacteria were added once to ensure enough food to support development. Early embryos were harvested from gravid adults. These embryos were either used directly to isolate embryonic PGCs or incubated for 12~16 hours in M9 solution until reaching the L1 stage for PGCs isolation. To isolate L1 PGCs from fed animals, the L1s were plated onto RNAi plates for additional 5 hours before processing for PGC isolation. For RNA-seq experiments described in Figure 1 and Figure 2, RNAi treatments were done at 25°C. For the rest of RNA-seq experiments, RNAi treatments were done at 20°C. See Supplemental Table S8 for sequencing library information.

To isolate PGCs from embryos, cell dissociation was performed as described in Strange et al. 2007 (Strange et al., 2007) with the following modifications: 1x106 embryos were treated in 500ul chitinase solution (4.2 unit of chitinase (Sigma # C6137) in 1ml of conditioned egg buffer). After chitinase treatment, embryos were collected by centrifugation at ~900g for 4 mins at 4°C; and resuspended in 500μl accumix-egg buffer solution for dissociation (Innovative Cell Techologies, AM105, 1:3 dilution ratio in egg buffer). In the final step, cells were resuspended in chilled egg buffer before sorting using BD FACSAriaII. 65,000~120,000 PGL-1::GFP PGCs were used for RNA isolation.

To isolate PGCs from L1 larvae, 400,000 to 500,000 packed L1s were used for cell dissociation as described in Zhang and Kuhn (Zhang and Kuhn, 2013)(www.wormbook.org/chapters/www_cellculture/cellculture.html#sec6-2) with the following modifications: starved and fed (for 5 hours) L1 were incubated with freshly thawed SDS-DTT solution for 2 min and 3min, respectively, with gentle agitation using a 1000μl pipette tip. Pronase treatment was performed using 150 μl of 15mg/ml pronase (Sigma P6911). Pronase treatment was stopped by adding 1000μl conditioned L-15medium and spin at 1600g for 6 min. Cells were resuspended in chilled egg buffer and washed three times to remove debris before sorting using BD FACSAriaII or Beckman Coulter MoFlo sorter. ~75,000 sorted cells were pelleted at 1600g for 5 mins, snap freezed and saved in −8003B1C for later RNAseq analysis.

## RNA extraction

RNA was extracted from sorted cells using TRIZOL. The aqueous phase was transferred to Zymo-SpinTM IC Column (Zymo research R1013) for concentration and DNase I treatment as described in manual. RNA quality was assayed by Agilent Bioanalyzer using Agilent RNA 6000 Pico Chip. All RNAs used for library preparation had RIN (RNA integrity number) >8.

### RNAseq library preparation and analysis

Three different RNA-seq library preparation methods were used for this study: SMART-seq, which uses poly-A selection (Figure 1 and 2), Nugen Ovation, which uses random priming (Figure S2), and Truseq combined with Ribozero to remove ribosomal RNAs (all other figures). The first two methods allow library construction from <10ng of total RNA, whereas the latter method requires >50ng total RNA. We compared SMART-seq and Truseq-Ribo zero performance on L1 PGCs isolated from wild-type and *nos-1(gv5)nos-2(RNAi)* and observed identical trends, with an overall higher number of miss-regulated genes identified with Truseq-Ribozero (Compare Figure 1 (SMART-seq) and Figure S1B-D (Truseq/Ribozero). For the experiment shown in Figure S2 where we compared RNA levels between embryonic PGCs and an oocyte library reference, we used Nugen Ovation libraries which avoids any bias due to poly-A selection while allowing library construction from < 3ng of RNA. For all experiments, control and experimental libraries were made using the same method. Table S5 contains lists of miss-regulated genes from analyses. Table S8 lists all the RNA-seq libraries used in this study and the corresponding figures.

SMART-seq libraries: libraries were made from 2ng of total RNA isolated from sorted PGCs from worms grown at 25°C using SMART-seq v4 Ultra Low input RNA kit (Clontech, Cat. No. 634888) followed by Low Input Library Prep Kit (Clontech, Cat. No. 634947). The cDNAs were then fragmented using Covaris AFA system at the Johns Hopkins University microarray core and cloned using the Low Input library prep Kit.

Nugen Ovation libraries: libraries were made from 3ng of total RNA isolated from sorted cells from worms grown at 25°C using Nugen Ovation system V2 (# 7102-08) followed by Nugen Ultralow library system.

TruSeq libraries: 50ng of total RNA isolated from sorted PGCs from L1 worms grown at 20°C; were subjected to Ribozero kit (illumina, MRZE706) to remove rRNA. Libraries were constructed using Truseq Library Prep Kit V2.

All cDNA libraries were sequenced using the Illumina Hiseq2000/2500 platform. Differential expression analysis was done using Tophat (V.2.0.8) and Cufflink (V 2.0.2). The command lines for Tuxedo suit are listed as below:

$ tophat2 -p 12 -g 1 --output-dir <*output>* --segment-length 20 --min-intron-length 10 --max-intron-length 25000 -G <*gene.gtf>* --transcriptome-index <Name.*fastq*>$ cuffdiff -p 12 -o <*output>* --compatible-hits-norm --upper-quartile-norm -b <genome.fa> <genes.gtf> <*tophat output1> <tophat output2>*

Gene set enrichment analysis for 4 different categories and correlation of gene expression were done using custom R scripts. Plots were drawn using R package and Prism software. In Figure 6F, the area-proportional Venn diagram was created using the VennDiagram R package. For comparisons shown in Figure S2A, oocyte transcriptome data was extracted from Stoeckius et al. 2014, and embryonic soma and germ cells expression profiles were from this study (Supplemental Table S8). Expression for each genes were log10 transformed, ranked and ordered. Correlations were plotted using custom R codes.

### ATACseq library preparation and analysis

ATACseq was performed as described in Buenrostro et al. 2015. Experimental pipelines was described as follows:

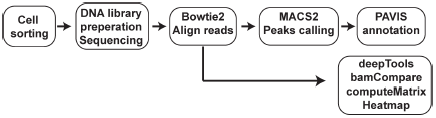

30,000 sorted L1 PGCs were washed with 60μl cold cell culture grade PBS once and spun at 2000g for 10min. Cell nuclei were isolated by resuspending cell pellets in cold lysis buffer (10mM Tris-Cl pH7.4, 10mM NaCl, 3mM MgCl2, 0.1% Igepal CA-630) followed by centrifugation at 3500g for 10 min at 40°C. The transposition reaction was performed with a 50μl reaction mixture (25μl TD, 2.5μl TDE, 22.5 ;Cl nuclease-free H2O. Illumina, Nextera DNA library preparation Kit FC-121-1030) at 37°C; for 30 min. Transposed DNA was purified using Qiagen MinElute kit and saved in −20°C. qPCR was used to determine appropriate PCR cycle number for PCR amplification as detailed in Buenrostro et al. 6-7 cycles of PCR amplification were used. Final cDNA libraries (150bp to 700bp) were selected using Agencourt AMPure beads (Beckman-Coulter A63880). Two biological samples for wild type and *nos-1(gv5)nos-2(RNAi)* were sequenced with Hiseq2500 platform.

Raw reads of two biological samples were first merged using *“cat”* command in Unix environment followed by bowtie2 (v2.1.0) alignment using *C. elegans* ce10 as the reference genome. MACS2 package was used to identify locus with nos-1/2 –dependent features (peaks). The *callpeak* function in MACS2 package is used as written here: macs2 callpeak -t < *nos-1/2* aligned reads.sam> -c < wild type aligned reads.sam> --nomodel --extsize 200 -f SAM -g ce --B -q 0.05. or macs2 callpeak -t < wild type aligned reads.sam> -c < *nos-1/2* aligned reads.sam >--nomodel --extsize 200 -f SAM -g ce --B -q 0.05.

In this function -c, mapped reads were used as a reference to identify *nos-1/2* dependent chromatin features. PAVIS (https://manticore.niehs.nih.gov/pavis2/) uses the output file NAME_summits.bed from MACS2 for peak annotation. Identification of genes with nos-1/2-dependent peaks at their upstream was extracted and gene IDs were cross-referenced with RNA-seq analysis in this study.To plot heatmap for ATAC-seq analysis, *bamCompare* and *computeMatrix* in deepTools package (http://deeptools.readthedocs.io/en/latest/) were used to visualize ATACseq profile of *nos-1/2*-dependent genes as shown in Figure 3A and S3A. Command lines were listed as below.*bamCompare -b1 <nos-1/2.bam> -b2<wild type.bam> -o <Name1.bw> --ratio ratio --normalizeUsingRPKM -ignore chrM -bs 10 -p max/2 computeMatrix reference-point --referencePoint TSS -b 2000 -a 2000 -R <nos-1/2-dependent_gene.bed> -S<Name1.bw> -o <Name2.gz> --sortUsing max --skipZeros -bs 10 -p 2 plotHeatmap -m <Name2.gz> --zMin 0 --colorList --heatmapHeight 20 --heatmapWidth 5 -out <heatmap.png>*

### Quantitative RT-PCR assay

To verify our analysis pipeline for RNAseq data, quantitative RT-PCR (qRT-PCR) reactions using sequencing libraries as templates were performed. The cDNA libraries were diluted to 1nM before performing qRT-PCR. Primers for qRT-PCR were listed in Supplemental Table S7. Enrichment of target mRNAs between wild type and *nos-1/2* was calculated using ΔΔCt with *tbb-2* expression then normalized to wild type control. Fold change were plotted and significance was calculated by paired t-test.

### Technical v biological replicates

Biological replicates refer to experiments performed on independently treated hermaphrodites (in the case of RNA-seq libraries, this refers to worms exposed to independent RNAi treatments followed by cell sorting and RNA extraction). All in vivo technical replicates refer to observations in the same strain from separate zygotes.

### Datasets

Datasets generated in this paper are available at GEO accession GSE100651 for ATAC-seq and GSE100652 for RNA-seq.

---

**Supplemental Figure S1.**
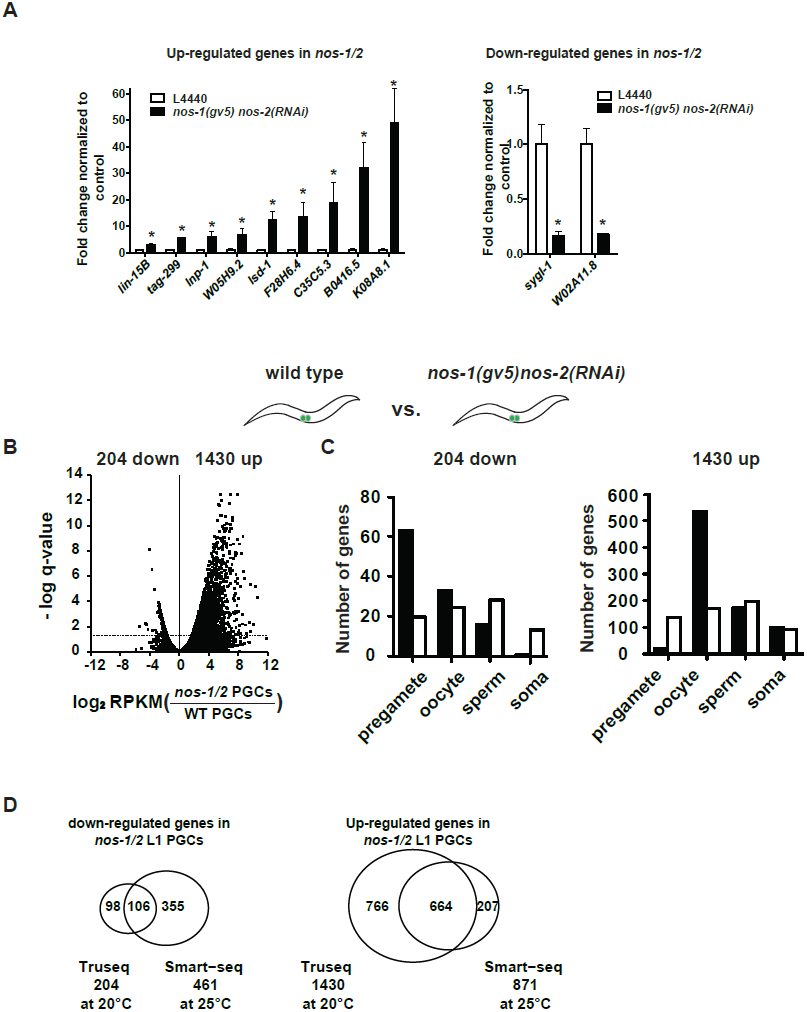
*nos-1nos-2* PGCs upregulate oogenic genes. (A) Bar graph showing results of quantitative RT-PCR of 11 genes significantly miss-expressed in *nos-1(gv5)nos-2(RNAi)* PGCs.^$^ (one asterisk: p < 0.05 using Student‘s t-test) (B-C) Transcriptome comparison between PGCs isolated from wild-type and *nos-1(gv5)nos-2(RNAi)* L1 larvae using Truseq libraries (see methods). (B) Volcano plot showing log2 fold-change of gene expression between PGCs isolated from wild-type and *nos-1(gv5)nos-2(RNAi)* L1 larvae. The numbers of genes that were significantly up- or down-regulated in *nos-1(gv5)nos-2(RNAi)* PGCs are indicated. Dashed lines mark the significance cutoff of q = 0.05 above which genes were counted as miss-expressed. (C) Bar graph showing expected and observed number of genes (Y axis) in different expression categories (X axis). (D) Venn diagrams showing overlapping genes between Tru-seq and SMART-seq analyses for genes miss-regulated in *nos-1(gv5)nos-2(RNAi)* compared to wild-type.

**Supplemental Figure S2.**
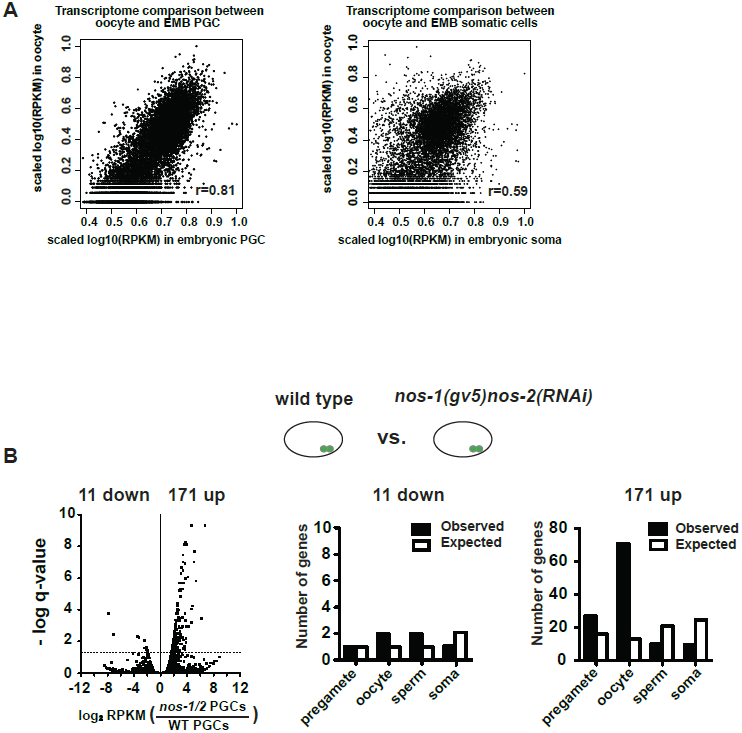
Maternal RNAs are maintained in early embryonic PGCs. (A) XY scatter plots showing correlation of gene expression between wild type embryonic PGCs (X axis) and oocyte transcriptome (Y axis) (Left) and correlation of gene expression between wild-type embryonic somatic blastomeres (X axis) and oocyte transcriptome (Y axis) (Right). High Pearson‘s correlation value was obtained for embryonic PGCs versus oocyte (R=0.81), but not for embryonic soma versus oocyte (R=0.59). For these analyses, information of oocyte transcriptome was obtained from Stoeckius et al., 2014. To compare two transcriptomes, level of gene expression (RPKM) was first subjected to log10 transformation and followed by internal scaling with the range of gene expression in each transcriptome. Plots were generated using R. (B) Transcriptome comparison between PGCs isolated from wild-type and *nos-1(gv5)nos-2(RNAi)* embryonic PGCs. Left: Volcano plot showing log2 fold-change of gene expression between PGCs isolated from *nos-1(gv5)nos-2(RNAi)* and wild-type embryonic PGCs. Middle and Right: Bar graph showing expected and observed number of genes in the different expression categories.

**Supplemental Figure S3.**
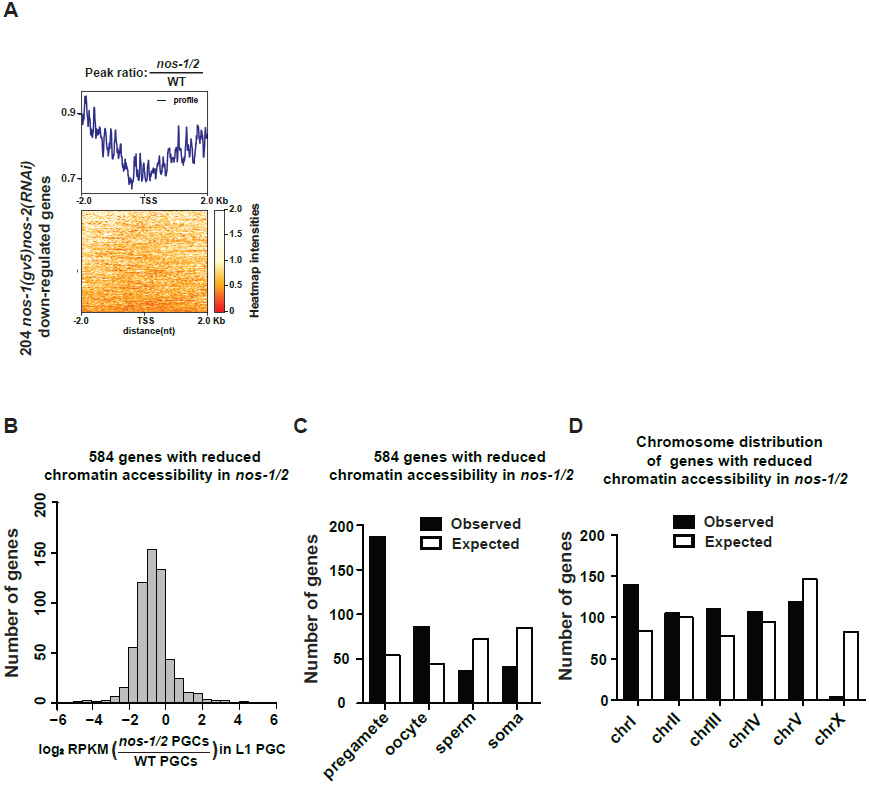
*nos-1(gv5)nos-2(RNAi)* PGCs fail to fully activate expression of pre-gamete genes. (A) Heat map showing ATAC-seq reads for 204 down-regulated genes in *nos-1(gv5)nos-2(RNAi)* compared to wild-type L1 PGCs. 4kb across transcription start site (TSS) were plotted on the heatmap. In the heatmap, darker color (red) indicates more reads (open chromatin) in wild-type PGCs. See materials and methods for detail description of data analysis. (B) Histogram showing the distribution of log2 fold change in gene expression between *nos-1(gv5)nos-2(RNAi)* and wild-type PGCs for 584 genes that acquired more repressive chromatin structure in *nos-1(gv5)nos-2(RNAi)* compared to wild-type L1 PGCs. Consistent with ATAC-seq denoting open/close chromatin features, genes with less open chromatin structure in *nos-1(gv5)nos-2(RNAi)* have decreased expression level compared to wild-type. See Table S2 for a list of 584 genes with more closed chromatin structure in *nos-1(gv5)nos-2(RNAi)* PGCs compared to wild-type PGCs. (C) Bar graph showing expected and observed number of genes acquired more repressive chromatin structure in *nos-1(gv5)nos-2(RNAi)* compared to wild-type L1 in four different expression categories. Consistent with previous RNA-seq analysis, *nos-1(gv5)nos-2(RNAi)* PGCs failed to activate pre-gamete genes. (D) Bar graph showing the chromosomal distribution of 584 genes that acquired more repressive chromatin structure in *nos-1(gv5)nos-2(RNAi)* compared to wild-type L1.

**Supplemental Figure S4.**
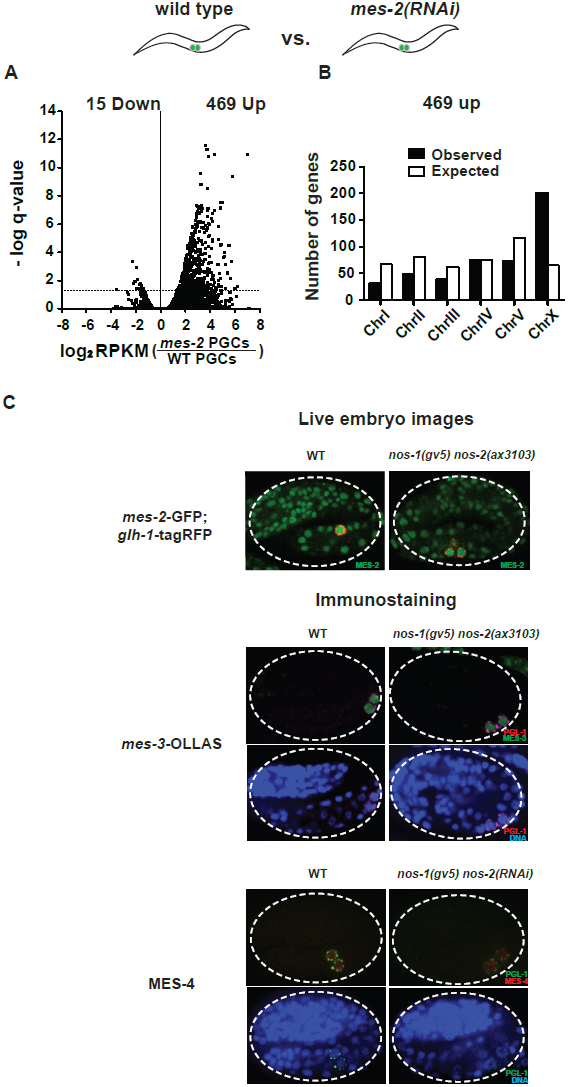
*mes-2* PGC transcriptomics and MES protein expression in in *nos-1nos-2* embryonic PGCs. Transcriptome comparison between PGCs isolated from wild-type and *mes-2(RNAi)* L1 larvae. (A) Volcano plot showing log2 fold change of gene expression between *mes-2(RNAi)* and wild-type L1 PGCs. The numbers of genes whose expression were up- or down-regulated in *mes-2(RNAi)* compared to wild-type L1 PGCs are indicated. Dashed lines mark the significance cutoff of q = 0.05 above which genes were counted as miss-expressed. (B) Bar graph showing chromosomal distribution of *mes-2(RNAi)* up-regulated genes. (C) Top: Photomicrograph of live embryo expressing GFP tagged MES-2 in wild-type and *nos-1(gv5)nos-2(ax3103)* embryos. Middle: Photomicrograph of fixed wild-type and *nos-1(gv5)nos-2(ax3103)* embryos expressing OLLAS tagged MES-3. Bottom: Photomicrograph of fixed wild-type and *nos-1(gv5)nos-2(RNAi)* embryos stained with anti-MES-4 antibody and K76 anti-PGL-1 antibody. Images of 2-fold+ stage embryos were taken.

**Supplemental Figure S5.**
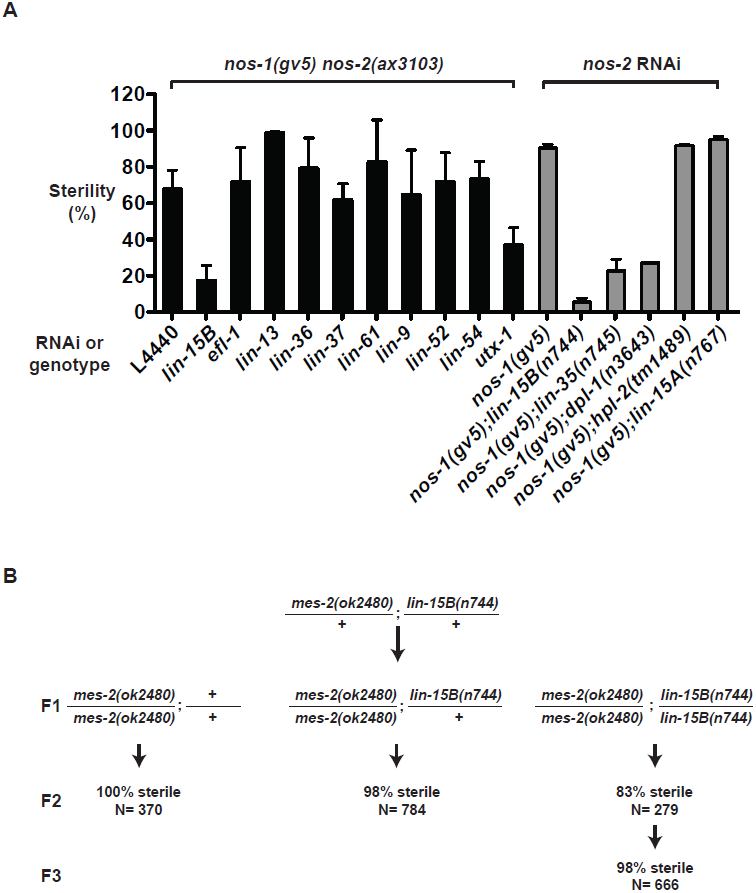
Suppression of *nos-1nos-2* sterility by *synMuvB* mutants. (A) Bar graphs showing percent sterility among worms of the indicated genotypes. Black bars: *nos-1(gv5)nos-2(ax3103)* mutant were fed with bacteria expressing dsRNA to genes indicated and the sterility of their progeny were scored. Gray bars: *nos-1(gv5)*; synMuvB double mutants were fed with bacteria expressing *nos-2* dsRNA and the sterility of their progenies were scored. Error bars report S.D. from ≥ 2 experiments. (B) Scheme for testing maternal effect sterility of *mes-2(ok2480); lin-15B(n744)* animals. First generation (F1) *mes-2(ok2480); lin-15B(n744)* animals were derived from *mes-2(ok2480)/+; lin-15B(n744)/+* hermaphrodites. The sterility of their progeny (F2) was scored. Loss of *lin-15B* suppressed *mes-2(vc2480)* maternal-effect sterility weakly for one generation (83% sterile F2). The percent sterility rose back up in the F3 generation (98%) and the line could not be maintained.

**Supplemental Figure S6.**
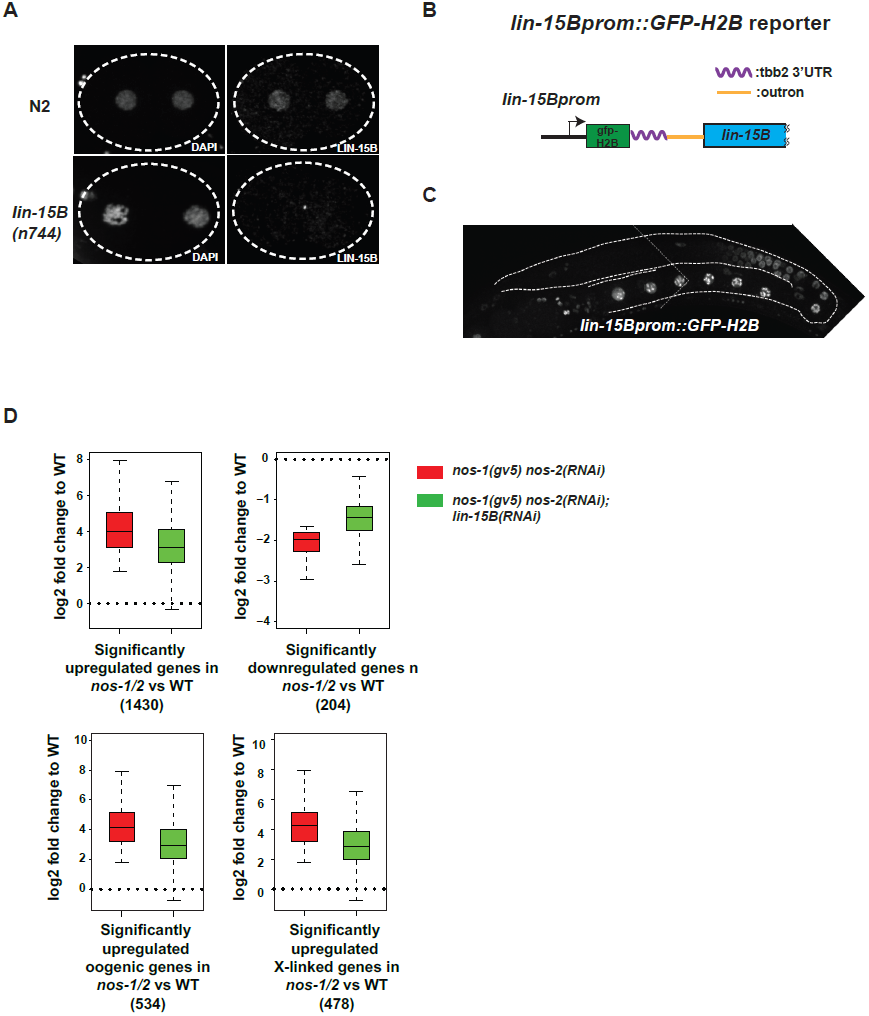
Loss of *lin-15B* mitigates gene expression changes in *nos-1nos-2* PGCs. (A) Photomicrograph of 2 cells wild-type and *lin-15B(n744)* embryos stained with anti-LIN-15B antibody. Anti-LIN-15B is specific as no nuclear signal was detected in *lin-15B(n744)* embryos. (B) Cartoon diagram showing the *lin-15B* transcriptional reporter. *GFP::Histone-H2B*::*tbb-2 3‘ UTR ::gpd-2/3 outron* was inserted at 5‘ end of *lin-15B* ORF in an operon configuration to preserve the function of endogenous *lin-15B*. (C) Photomicrograph of adult hermaphrodite expressing the *lin-15B* transcriptional reporter. Germline is outlined. Expression in the *lin-15B* promoter reporter begins in late pachytene germ cells committed to oogenesis. (D) Box and Whisker plot showing log2 fold change compared to wild-type of different categories as depicted under each plot. Each box extends from the 25th to the 75th percentile, with the median indicated by the horizontal line; whiskers extend from the 2.5th to the 97.5th percentiles. The level of miss-regulation of gene expression is reduced in *nos-1(gv5)nos-2(RNAi); lin-15B(RNAi)* PGCs.

**Supplemental Figure S7.**
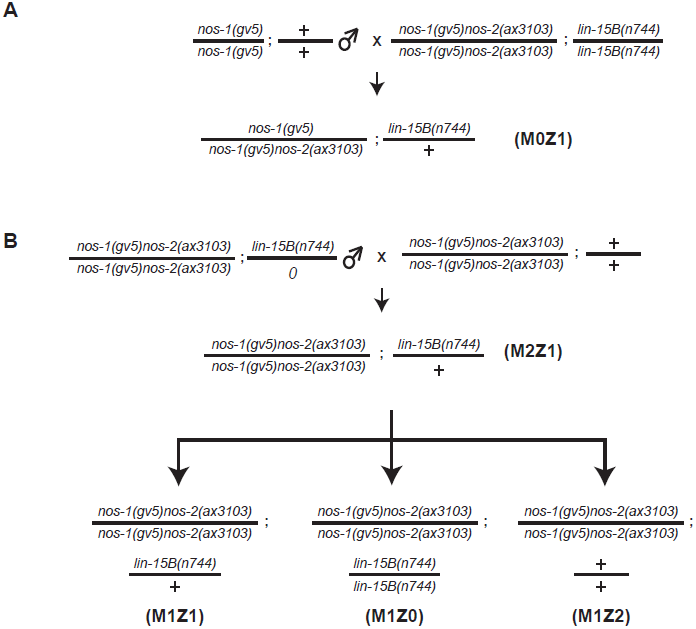
Assay for maternal and zygotic contribution of *lin-15B* in *nos-1nos-2* sterility. (A-B) Mating schemes to characterize the maternal and zygotic contribution of *lin-15B* in *nos-1(gv5)nos-2(ax3103)*. *lin-15B* genotypes were determined by Sanger sequencing (See Table S7 for PCR/sequencing oligos).

